# Ivory-billed Woodpecker and Blue Jay kent comparisons at spectrogram level reveal differences

**DOI:** 10.1101/2024.10.25.620249

**Authors:** John D Williams

**Affiliations:** Mission Ivorybill

## Abstract

A representative audio file for Blue Jay (Cyanocitta cristata), that is thought to be most similar to suspected kent sounds of the Ivory-billed Woodpecker (Campephilus principalis), was examined at the spectrogram level. A total of 136 other Blue Jay files were looked at and compared to kents (n>200) from six Ivory-billed Woodpecker expeditions. At this precision, differences are seen such that these two species cannot be mistaken for each other.

## Introduction

Bird sounds can be identified and assigned to a species. The continuing search for evidence that the Ivory-billed Woodpecker exists includes the acquisition of audio recordings. This has been accomplished numerous times since 1935, with an increase since 2000 [1]. The proper examination and identification of potential sources, and elimination of other possibilities, can potentially be considered proof that this species is not extinct.

Blue Jays are notorious mimics and also have a wide repertoire of sounds. They can make kentlike sounds that people believe instead are from the Ivory-billed Woodpecker. This paper began with a request to examine a Blue Jay (BLJA) sound file, which to the ear sounds like audio recorded by Ivory-billed Woodpecker (IBWO) expeditions.

## Methods

The program Sonic Visualizer [2] was used to create a spectrogram of the calls and then compare it with suspected IBWO recordings and spectrograms made from them. All spectrograms were adjusted to be the same X and Y range, where X is about 4000 Hz (Hertz which measures tone) and Y is about 2.2 seconds. When spectrograms are examined with this precision, details emerge that can show similarities or differences.

## Results

The Blue Jay sound examined was from the Macaulay Library BLJA file (ML169755) [3]. Below is a spectrogram from this file at the 40 second mark. Some professionals believe this particular section of the recording to be the Blue Jay example most similar with IBWO:

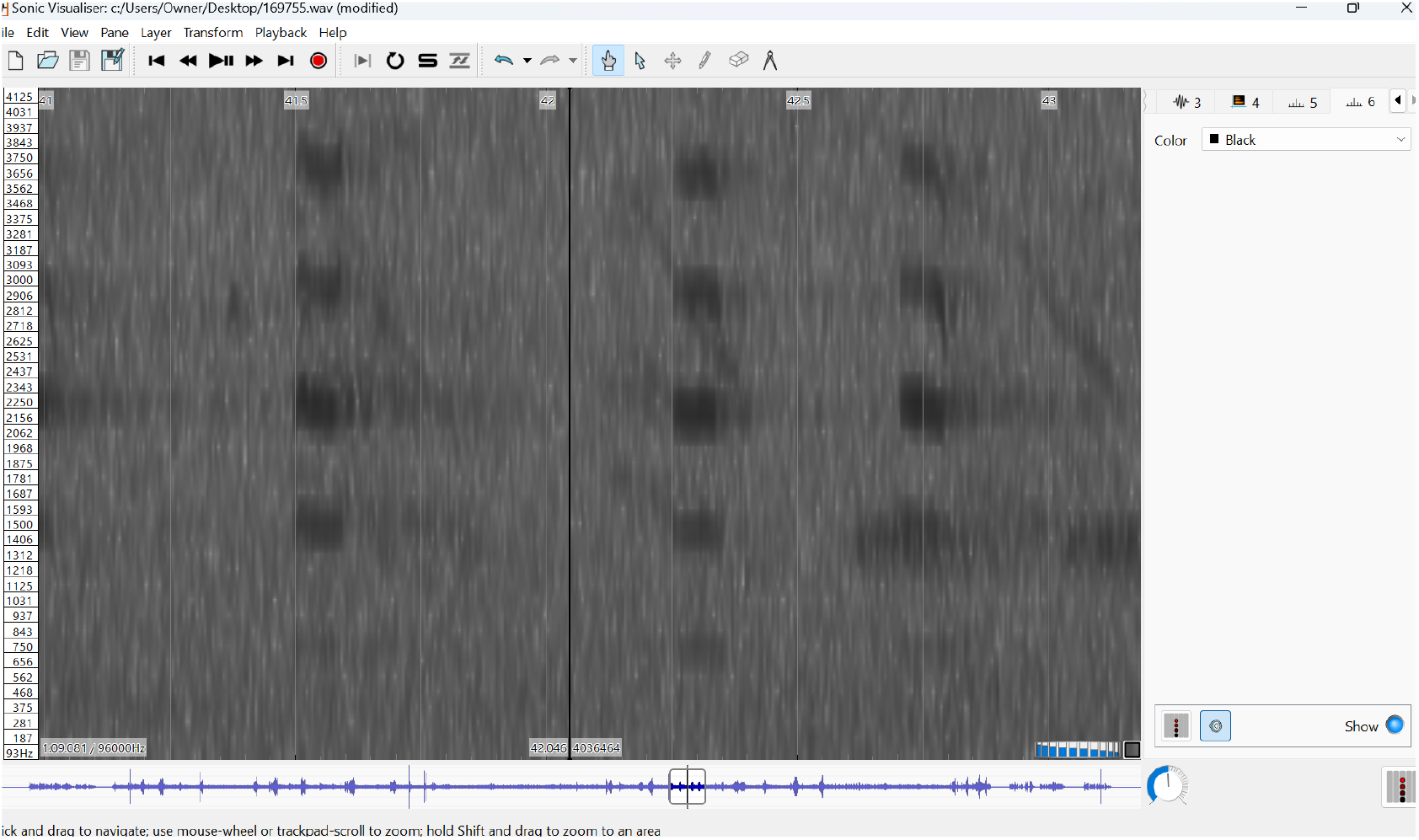
Image 1 [3]

Note the blocky, squarish pattern. All the kent-like sounds on this recording have this look. They are short, approximately 0.1 second each. They sound to the ear around 734 Hz, with one at 693 and another around 719 Hz. There are also some sounds that do not show as flat harmonic lines, but as lines with slopes. These are around 743 Hz, have a harsh quality and are what many people call typical BLJA sounds.

An additional 136 audio files of BLJA were examined from the Macaulay Library [4]. These either do not look similar, do not sound similar, or both, to suspected IBWO recordings. None of the 136 files examined were close to 0.3 seconds, which occur repeatedly in what are thought to be IBWO kent calls, and none of these 136 BLJA files were at 660 Hz as were suspected IBWOs recorded by Cornell in Arkansas, and not near 587 Hz as recorded by numerous other IBWO-focused expeditions [5].

Below are representative spectrograms from IBWO expedition sources. The kent sounds appear as horizontal, ladder-like marks. There are clear differences from the BLJA ML169755 file:

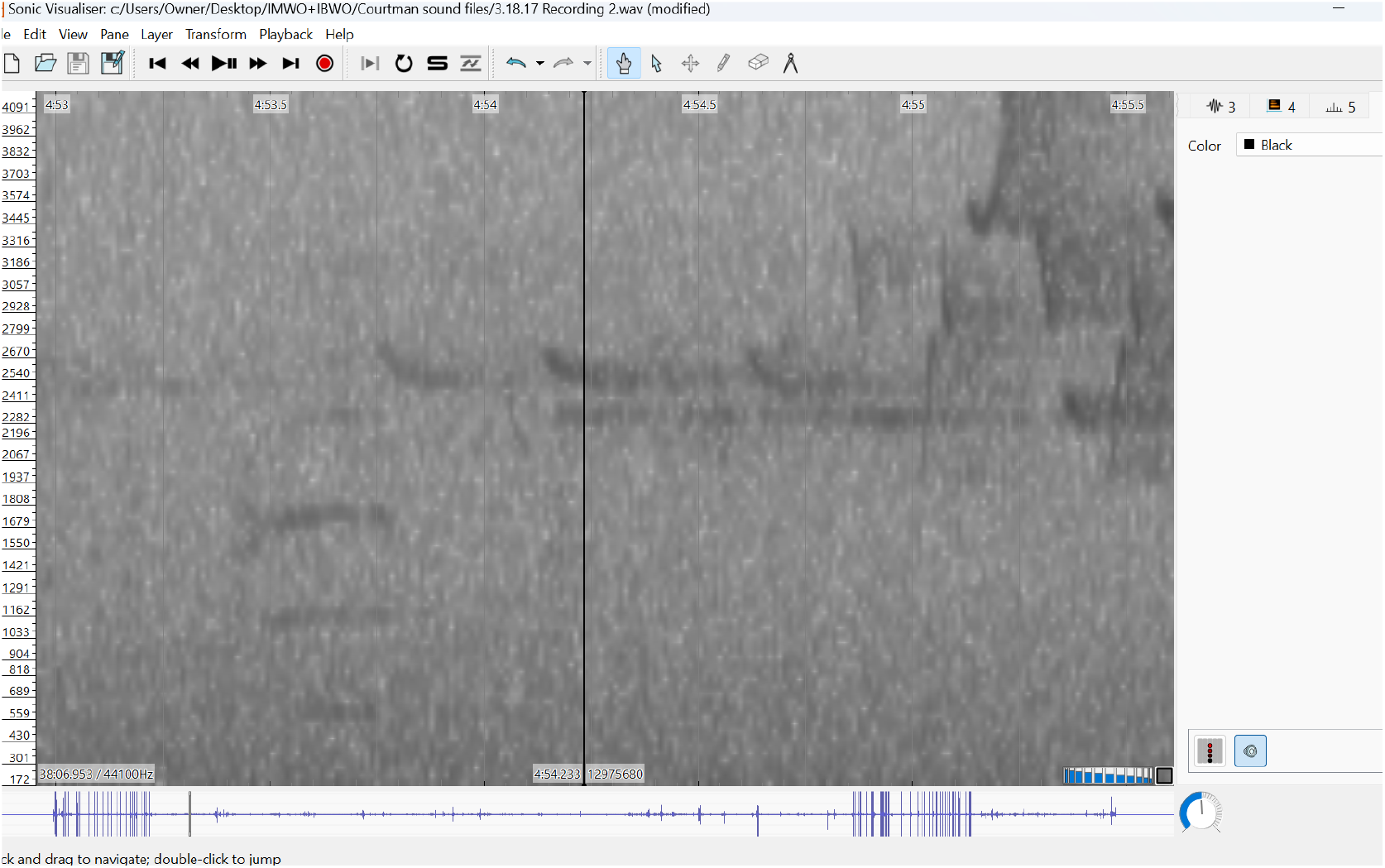
Image 2, Matt Courtman [6]

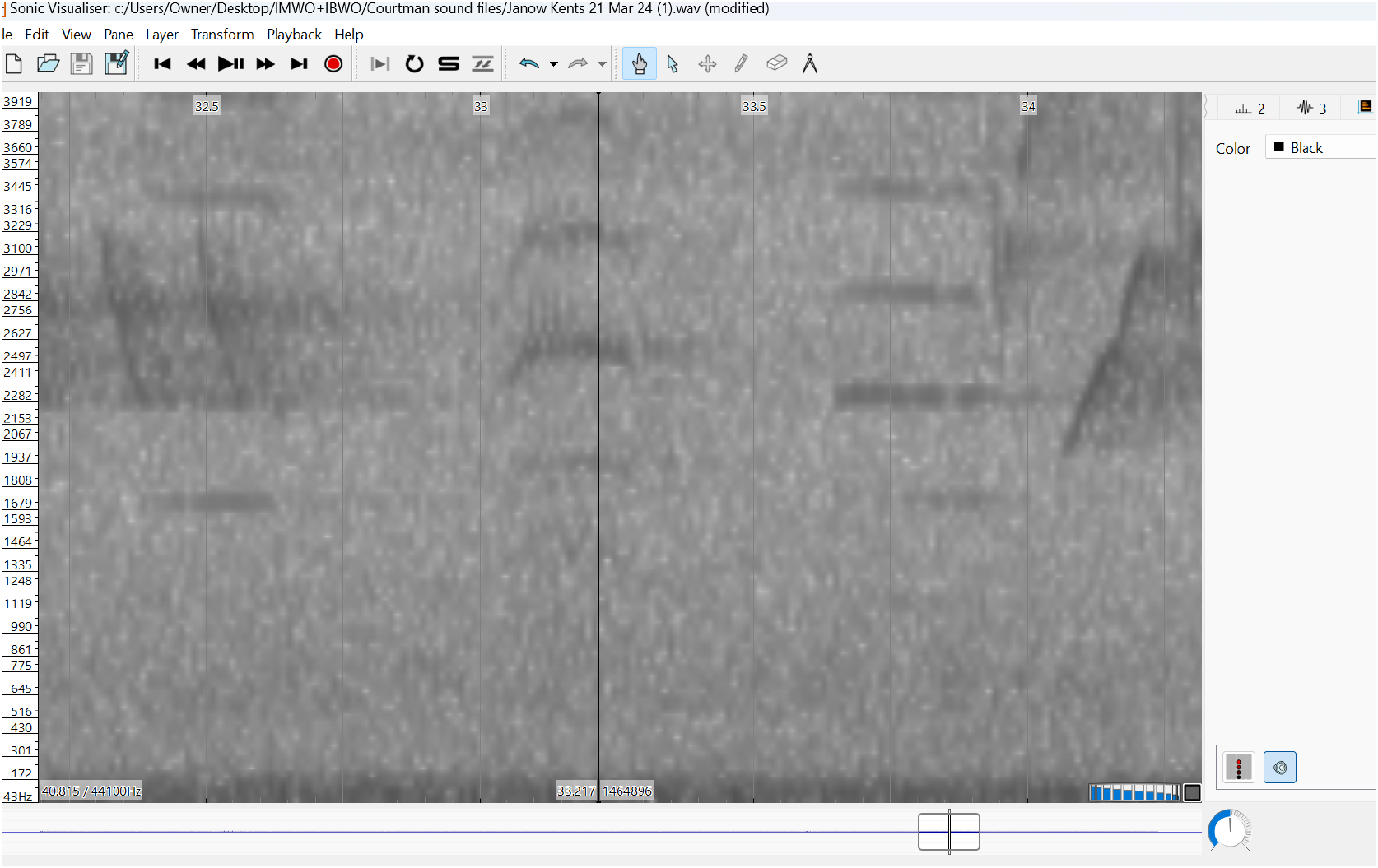
Image 3, Peter Janow [7]

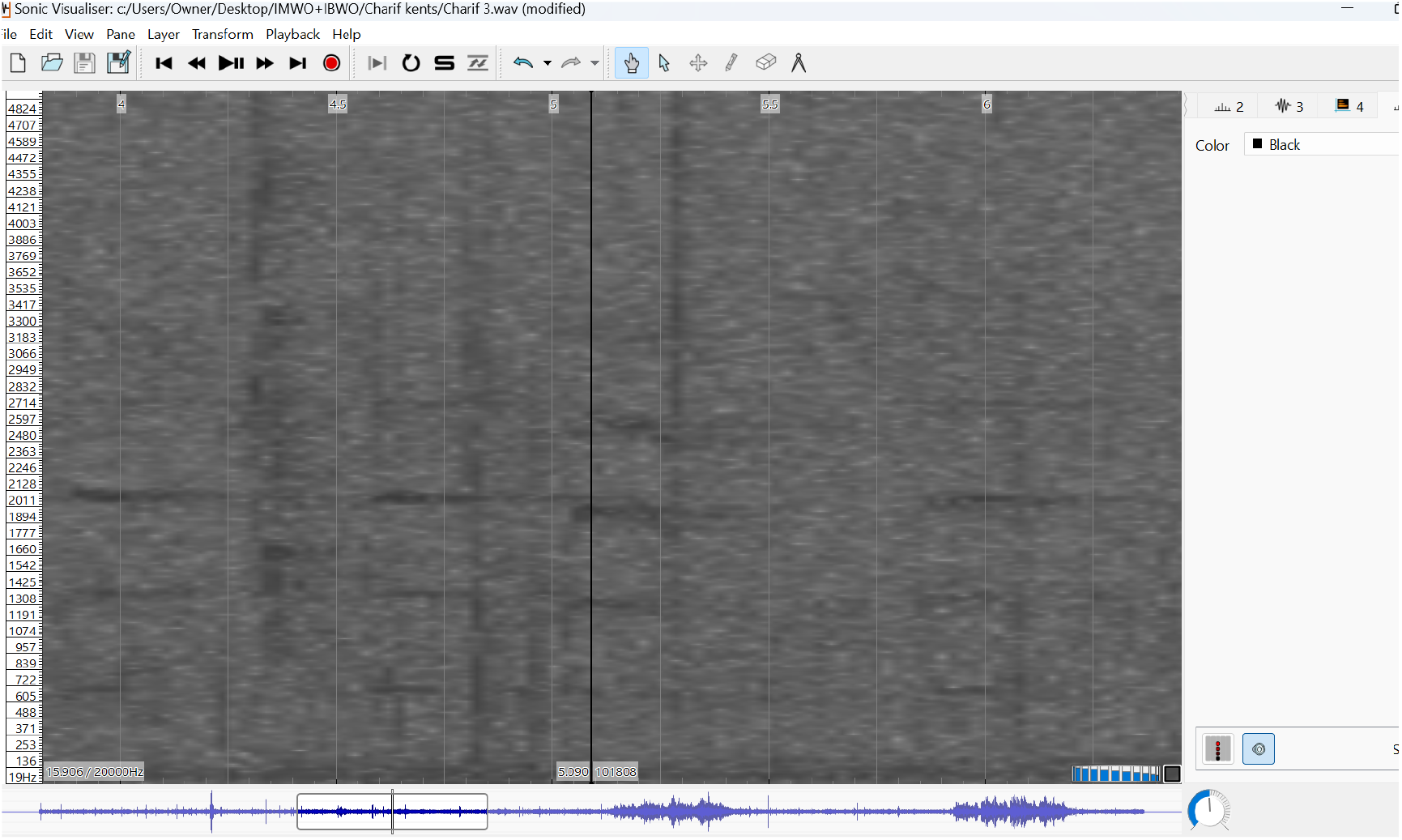
Image 4, Russ Charif [8]

Dan Mennill’s numerous recordings of suspected IBWO kents (n= 208), from the Geoff Hill 2005-6 expedition [9], look similar, as do the recordings made by John Dennis in 1968 [10].

Almost all of these suspected IBWO kent calls are longer at around 0.3 seconds, sound lower, often at 587 Hz, and the spectrogram lines are narrower, showing a more pure tone with less dissonance. Recordings obtained by Courtman, Janow, Charif, Mennill, Dennis, and other unpublished recordings are quite similar in spectrogram structure, so the examples above are representative. See [5] for further explanation with audio samples.

## Conclusion

Because of different morphology and functional anatomy, Blue Jays and Ivory-billed Woodpeckers are going to make different quality sounds if examined at sufficient precision. These differences become visible on a spectrogram. The closer one looks, the more apparent it is that the BLJA “kent” call is different. It is shorter at about 0.1 second, and each spectrogram mark is more vertical. In other words, each of the harmonic lines covers a greater Hz or tonal range. This equals non-harmonic dissonance.

The sounds that are thought to be from the Ivory-billed Woodpecker– obtained by professional expeditions which included sightings, and in some cases imagery and videos– are of 0.3 seconds in length, around 587 to 660 Hz, and have narrow lines which imply pure tone.

A previous and similar study by Hardy [11] arrived at a similar conclusion using a different method. He examined Blue Jay spectrograms that appeared similar to John Dennis’ suspected IBWO recording [10], but found the audible Hz to be higher in each case. He notes that the Blue Jay sounds are “metallic”-- this quality has never been stated for suspected IBWO kents. Metallic sounds imply harshness and a higher tone.

All suspected Ivory-billed Woodpecker recordings, including Dennis’ in 1968, and from numerous sources since 2000, also agree at spectrogram level with the only proven IBWO recording from the 1935 Cornell expedition, where birds were filmed at their nest [12].

Geoff Hill, on his web page Ivory-billed Woodpeckers in the Florida Panhandle, also addresses the Blue Jay question:

“One could perhaps dismiss our recorded kent calls as atypical vocalizations of Blue Jays although Blue Jays were absent from all but the edge of our study site throughout the winter and members of our search team have yet to hear Blue Jays make sounds like our putative kent calls. Moreover, typical Blue Jay sounds are never recorded in association with these kent calls. Because jays are vocal birds, it is unlikely that our kent recordings can be attributed to Blue Jays silently approaching a microphone, giving one or a few kent notes, and then slipping silently away… What are the chances that rare Blue Jay kent calls and rare pileated double knocks would occur together in the same spot on ten separate days of recording? It seems to us that a knocking and vocalizing Ivory-billed Woodpecker as the source of the kents and double knocks is the simpler explanation, especially when the location of the sound recording is where 13 sightings were made and there are many scaled trees and large cavities.” [13]

Those who become interested in the Ivory-billed Woodpecker story, its intrigue, legend, and ideas of proof, encounter the critique that the sounds recorded for it are actually from the much more common, and excellent mimic, Blue Jay. This paper serves to show that this idea is improbable, perhaps impossible, and that the notion of “extraordinary claims require extraordinary evidence” is met with the use of precision spectrogram analysis. The result favors the continuing existence of the Ivory-billed Woodpecker.

The author declares no competing interests. Contact at

